# NucleoMap: a computational tool for identifying nucleosomes in ultra-high resolution contact maps

**DOI:** 10.1101/2021.11.15.468663

**Authors:** Yuanhao Huang, Bingjiang Wang, Jie Liu

## Abstract

Although poorly positioned nucleosomes are ubiquitous in the prokaryote genome, they are difficult to identify with existing nucleosome identification methods. Recently available enhanced high-throughput chromatin conformation capture techniques such as Micro-C, DNase Hi-C, and Hi-CO characterize nucleosome-level chromatin proximity, probing the positions of mono-nucleosomes and the spacing between nucleosome pairs at the same time, enabling profiling of nucleosomes in poorly positioned regions. Here we develop a novel computational approach, NucleoMap, to identify nucleosome positioning from ultra-high resolution chromatin contact maps. By integrating nucleosome binding preferences, read density, and pairing information, NucleoMap precisely locates nucleosomes in both eukaryotic and prokaryotic genomes and outperforms existing nucleosome identification methods in precision and recall. We rigorously characterize genome-wide association in eukaryotes between the spatial organization of mono-nucleosomes and their corresponding histone modifications, protein binding activities, and higher-order chromatin functions. We also predict two tetra-nucleosome folding structures in human embryonic stem cells using machine learning methods and analysis their distribution at different structural and functional regions. Based on the identified nucleosomes, nucleosome contact maps are constructed, reflecting the inter-nucleosome distances and preserving the original data’s contact distance profile.

## 2 INTRODUCTION

Nucleosomes are the conservative building blocks of the hierarchical chromatin structure, on which higher-order structures are formed.^1^ Genome-wide approaches revealed that nucleosomes are regularly spaced and organized into arrays in different species, but their positioning levels may vary.^2^ In yeast, almost all nucleosomes are well-positioned, meaning that they occupy the same location of the genome in most cells of a population.^3^ However, even though the space-regularity of nucleosome arrays are preserved, most of the nucleosomes are poorly positioned in animals and plant genomes.^4, 5,6^ Nucleosome positioning level is closely related to chromatin functions from two aspects. First, it has been shown that changes in the space-regularity and the nucleosome positioning level directly correlate with gene activity in eukaryotes^7, 8, 3, 9,10^ and prokaryotes.^11, 12,5^ Second, hi-stone modifications and non-histone binding proteins, which indirectly participate in the gene regulation, also affect nucleosome positioning.^13^ Despite their close relationship with the chromatin function, the underlying nucleosome distributions have not yet been comprehensively studied in eukaryote genomes due to the ubiquitous poorly-positioned nucleosomes.^14,6^

Current nucleosome identification methods rely on the peak patterns in MNase-seq,^15, 16, 17,18^ ChIP-seq,^19,20^ or ATAC-seq.^21,22^ However, poorly positioned nucleosomes yield broad and ill-defined peaks in the data,^14,4^ increasing the difficulties of locating them using current computational methods. While the aforementioned sequencing techniques capture only the occupancy of mono-nucleosomes along the genome, identifying poorly positioned nucleosomes requires including their space-regularity, which remains unchanged even in poorly positioned regions. Therefore, to understand the nucleosome distribution in eukaryotic genomes, especially in poorly positioned regions, it is necessary to integrate additional information from other data sources.

Recently available high-throughput enhanced chromatin conformation capture (Hi-C) techniques such as Micro-C,^23,24^ DNase Hi-C^25^ and Hi-CO^26^ provide information of nucleosome-level chromatin proximity. These ultra-high resolution chromatin contact map data capture both *mono-nucleosomes’ positions* characterized by the read alignments and the *spacing between nucleosome pairs* characterized by contact distances. Integrating nucleosome positioning and spacing information enables identifying nucleosomes in poorly positioned regions. With the increasing availability of the data, identifying nucleosomes from ultra-high resolution chromatin contact maps becomes meaningful. However, no computational approach has been specifically designed to call nucleosome positions from ultra-high resolution chromatin contact maps to our best knowledge.

In this work, we present NucleoMap, a nucleosome identification approach from ultra-high resolution chromatin contact maps. By integrating genomic sequence specificity, read density, and pairing information, NucleoMap precisely locates nucleosomes in both eukaryotic and prokaryotic genomes and outperforms existing nucleosome identification methods in precision and recall. We rigorously characterize genome-wide association in eukaryotes between the spatial organization of mono-nucleosomes and their corresponding histone modifications, protein binding activities, and higher-order chromatin functions. For the first time, we predict the distribution of two tetra-nucleosome folding motifs, *α*-tetrahedron and *β*-rhombus, in human embryonic stem cells. The association between preferences on folding motifs and genome structure is investigated. Based on the identified nucleosomes, nucleosome contact maps are constructed, which preserve the inter-nucleosome distances. In this way, nucleosome contact maps capture the original contact distance profile, making them more concentrated and more interpretable than the traditional fixed-bin-based contact maps.

## 3 RESULTS

### 3.1 NucleoMap algorithm

Most existing methods locate nucleosomes from aligned reads via a binding site model.^15, 16, 17, 18,19^ These methods use peak-calling approaches to identify genomic regions with enriched reads or normalized occupancy signals. In this way, nucleosome positions are solely determined by the peak patterns in the read density involving multiple nucleosomes. As a result, these models are not sensitive at identifying poorly positioned nucleosomes whose peaks are broad or ill-defined. To overcome the limitation, we develop an approach called NucleoMap. NucleoMap identifies nucleosomes at different positioning levels from ultra-high resolution chromatin contact maps, including Micro-C,^23,24^ DNase Hi-C^25^ and Hi-CO.^26^

Unlike traditional peak calling methods, NucleoMap uses a parametric model to separate the read density into multiple small distributions, each representing the positioning of an individual nucleosome. In particular, every nucleosome’s position is explicitly characterized by a Gaussian distribution, and NucleoMap identifies them by maximizing a likelihood function integrating aligned reads, nucleosome binding preferences, and pairing information in chromatin contact maps (Fig. 1).

**Figure 1:**
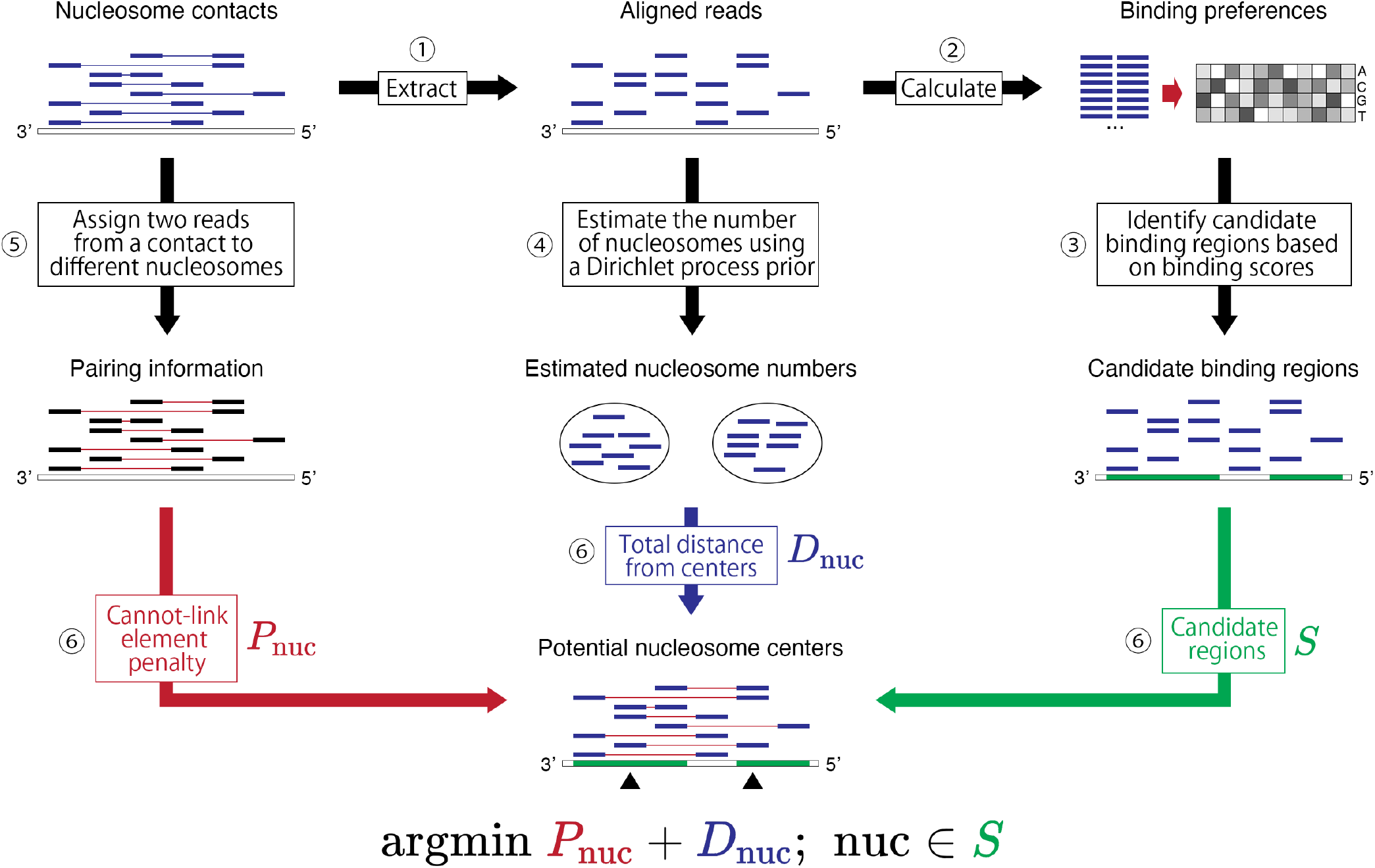
Workflow of the NucleoMap model. NucleoMap locates nucleosome centers with the following steps: **1.** Extract aligned reads from ultra-high resolution chromatin contact maps. **2.** Calculate nucleosome-binding preferences from the contact sequences. **3.** Identify candidate nucleosome-binding regions with the binding preference in the previous step. **4.** Estimate the number of nucleosomes using a Dirichlet prior. **5.** Extract the pairing information of reads from ultra-high resolution chromatin contact maps. Two ends from the same contact will be assigned to different nucleosomes. **6.** Calculate nucleosome centers in candidate binding regions by integrating the read positions and pairing information.

The first piece and the most direct information is the aligned read density, which is also used by traditional peak-calling approaches (Fig. 1 step 4). Since a read in a separate read density indicates a mono-nucleosome from an individual cell, NucleoMap defines the expected nucleosome position of a separate read density as its mean position, and the expected nucleosome positions within separate read densities are identified following the *k*-means paradigm. Due to the unknown number of nucleosomes in a region, NucleoMap first adaptively identifies *k* using a Dirichlet process (DP) prior. After that, reads are assigned to their nearest nucleosomes according to their distances to the density centers (Fig. 1 step 6).

The second piece of relevant information comes from nucleosome binding preference reflected by the nucleosome binding motifs (Fig. 1 step 3). It is known that nucleosomes are enriched for particular DNA sequence motifs on the nucleosomal DNA, most notably ~10bp periodic occurrences of AA/AT/TA/TT 2-mers.^27, 28,29^ NucleoMap models the AA/AT/TA/TT dinucleotide motifs using a position weight matrix (PWM) and calculates a motif-based nucleosome-binding score along the genome. By integrating the binding score as a part of our likelihood function, NucleoMap considers the sequence specificity in nucleosome identification (Fig. 1 step 6).

As the third piece of information, the pairing information in contact maps is utilized by NucleoMap to facilitate the separation of neighboring nucleosomes (Fig. 1 step 5). Because each nucleosome can be profiled at most once in individual cells during the sequencing procedure, the contacts carry hidden information that their two ends do not belong to the same nucleosome. In NucleoMap, the read pairs served as cannot-link elements are considered in the read assignment step, providing additional information to separate closely located or poorly positioned nucleosomes (Fig. 1 step 6).

Using a likelihood function integrating the three aforementioned types of information, NucleoMap is optimized by a *k*-means optimization procedure. In the end, the reads are separated into different clusters representing mono-nucleosomes, while the number of nucleosomes *k* is automatically learned using a hyperparameter *λ. λ* controls the fuzziness threshold of nucleosome calling.

### 3.2 NucleoMap accurately locates well-positioned and poorly-positioned nucleosomes

To directly evaluate the performance achieved by our method, we first compare the precision and recall of different nucleosome callers in yeast,^15, 20, 16,18^ where the positions of experimentally confirmed nucleosomes are available.^29^ Using these experimentally confirmed nucleosomes as the ground truth, the evaluation criteria are calculated as follows. First, the distance between a nucleosome position identified by the caller and its nearest experimentally confirmed nucleosome *d* is calculated. Then, nucleosomes with *d* shorter than a certain threshold are considered to be true-positive, meaning that they are validated by the experiment. In the end, the precision and recall are calculated for every method under different distance thresholds. We have the following observations. First, NucleoMap has the highest recall when distance threshold *d* is small, and it achieves the second highest recall at *d* =100bp (Fig. 2A). Compared with baseline methods, a significantly larger number of ground truth nucleosomes are identified by NucleoMap with *d* ≤80bp, suggesting its outstanding sensitivity in accurately identifying nucleosomes. At *d* =100bp, almost all ground truth nucleosomes are recovered by NucleoMap and DANPOS2, which recognize 1,554 and 1,622 out of 1,716 ground truth nucleosomes respectively. In comparison, only 1,162 ground truth nucleosomes are identified by NucTools, 417 are identified by NOrMAL and 346 are identified by Nseq. Second, NucleoMap also has the highest precision at every distance threshold d. Compared with baseline methods, the ground truth nucleosomes are composed of a higher ratio in nucleosomes identified by NucleoMap, suggesting a lower false-positive rate achieved by our method (Fig. 2B). Almost all (95.2%) nucleosomes identified by NucleoMap are ground truth nucleosomes at *d* =100bp. Ground truth nucleosomes identified by NOrMAL also account for a high proportion (95.1%), followed by NucTools (93.0%), DANPOS2 (88.9%), and Nseq (73.8%). Therefore, NucleoMap outperforms baseline models in both precision and recall.

**Figure 2:**
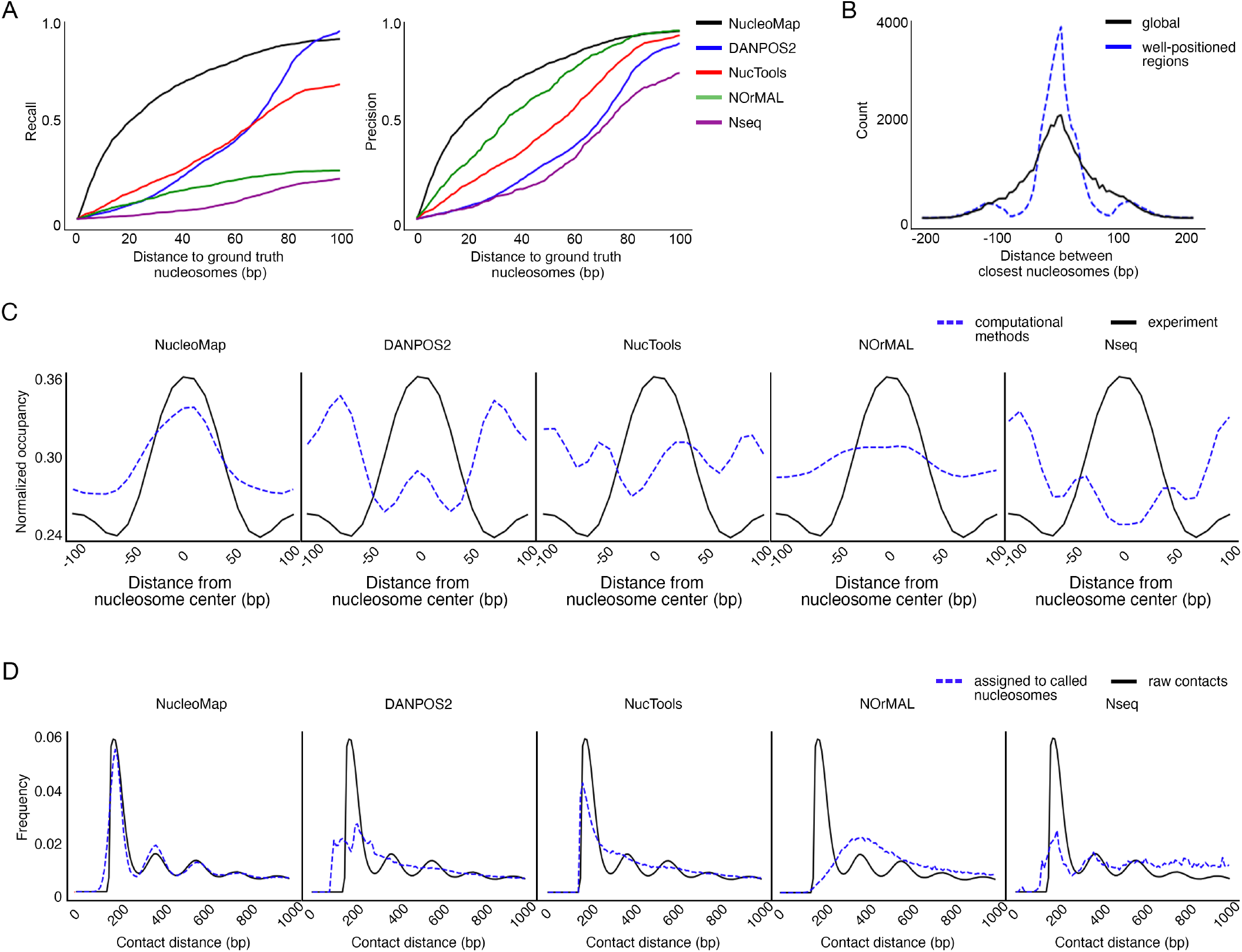
NucleoMap outperforms baseline methods in yeast and hESC. **A.** (left panel) Recall of nucleosomes identified by different approaches against the corresponding distance thresholds in yeast chrIII. The recall is calculated by n(True-positive nucleosomes)/n(Ground truth nucleosomes). Here “true-positive” nucleosomes refer to nucleosomes located within certain distance thresholds from a “ground truth” nucleosome, while the “ground truth” nucleosomes are experimentally confirmed nucleosomes. Smaller distance thresholds correspond to more accurate nucleosome locations, while higher recall corresponds to more ground truth nucleosomes identified by the methods. Therefore, the under area under the curves represents the sensitivity of the corresponding methods in identifying nucleosomes (right panel). The precision of nucleosomes identified by different approaches against the corresponding distance thresholds in yeast chrIII. Precision is calculated by n(True-positive nucleosomes)/n(identified nucleosomes). Higher consensus nucleosome ratio represents fewer “false-positive” nucleosomes identified by the methods, and thus the under area under the curves represent the nucleosome identifying specificity of the corresponding methods **B.** Distance between nucleosomes identified by NucleoMap (NucleoMap nucleosomes) and their nearest nucleosomes identified by DANPOS2 (DANPOS2 nucleosomes) in different regions. Compared with random samples collected from the whole genome, NucleoMap nucleosomes are much closer to the nearest DANPOS2 nucleosomes in well-positioned regions, showing the consistency in well-positioned regions across the two methods. **C.** Occupancy profile of experimentally identified nucleosomes (black) and nucleosomes identified by computational methods (blue) in yeast chrIII. The occupancy profile quantifies the average fraction of well-positioned nucleosomes within a ± 100bp window from the nucleosome centers. A similar profile of computationally identified nucleosomes with experimentally identified nucleosomes suggests accurate nucleosome identification. **D.** Occurring probabilities of short-range raw contacts (black) and contacts assigned to nucleosome centers called by computational methods (blue) against the genomic distance in hESC chr21. The peak patterns indicate contacts formed by neighboring nucleosomes and the distance between neighboring peaks represents the genome-wide nucleosome repeating length (NRL). Similar occurring probabilities between raw contacts and contacts assigned to nucleosome centers suggest accurate nucleosome identification.

To further prove that our method is able to identify both well-positioned and poorly-positioned nucleosomes, we compare the nucleosomes identified by NucleoMap and DANPOS2 in known well-positioned regions and random regions. In known well-positioned regions such as promoters, insulators, and enhancers, nucleosomes identified by NucleoMap are significantly closer (~ 50%) to their closest neighbors identified by DANPOS2, compared with random regions. The average distance between nucleosomes identified by NucleoMap and their closest neighbors identified by DANPOS2 in well-positioned regions is ~ 20bp. (Fig. 2B) This result suggests that NucleoMap performs at least as good as, if not better than, existing nucleosome calling methods in well-positioned regions.

To demonstrate the interpretability of our method, we then compare the nucleosome occupancy profile from the callers with experimentally identified nucleosomes in yeast. Nucleosome occupancy is a measure quantifying the fraction of occurring nucleosomes at a given position.^14^ Since most nucleosomes in yeast are well-positioned, a peak pattern in nucleosome occupancy is expected at yeast nucleosome centers. Indeed, experimentally identified nucleosomes in chrIII have a clear and symmetric peak-shape occupancy profile (Fig. 2C). This profile is used to validate the computationally identified nucleosomes.^29^ The average nucleosome occupancy of nucleosomes called by different methods is compared with the 1,716 ground truth nucleosomes in chrIII. Symmetric nucleosome occupancy profiles are observed at nucleosomes identified by NucleoMap, DANPOS2, NOrMAL, and Nseq, and occupancy peaks are observed in nucleosomes identified by NucleoMap, DANPOS2, and NOrMAL. Among the compared methods, NucleoMap identified nucleosomes have the most similar nucleosome occupancy profile with the occupancy profile of experimentally identified nucleosomes, suggesting that NucleoMap outperforms baseline methods.

Finally, to examine the overall performance of our method in eukaryotic genomes, we compare the recovered contact profile from the callers with the original contact profile in hESC Micro-C data. The contact profile of a contact map illustrates the contact frequencies at different contact distances. This profile reflects the real nucleosome spacing in the genome, providing a baseline for result validation. In order to compare the spacing of nucleosomes identified by callers with this baseline, we calculate their recovered contact profiles in two steps. First, two ends of a read are assigned to their nearest called nucleosomes, forming a recovered contact, and the distance between the assigned nucleosome pair is considered as the recovered contact distance. After that, the recovered contact profile is built using these recovered contacts, illustrating the spacing between computationally identified nucleosomes. Finally, the recovered contact profile is compared to the original contact profile. If the nucleosome spacing is consistent between the nucleosomes identified by callers and the underlying ground truth nucleosomes in the data, the two profiles are similar with each other. Compared with the nucleosomes called by baseline methods, nucleosomes identified by NucleoMap produce a more similar contact distance profile with the original data (Fig. 2D). This result implies that NucleoMap achieves high accuracy in identifying nucleosomes in prokaryotes.

### 3.3 Nucleosome positioning level and spatial organization reflect patterns of histone modification and genome functions

It has been discovered that nucleosome positioning reflects the genome functions in different regions because the nucleosomes are directly decorated, composed, or impeded by specific histone variants and regulatory proteins.^11,30^ To further validate the identified nucleosomes, as well as to evaluate the connection between nucleosome spatial distribution and genome functions, nucleosome positioning levels and local nucleosome organization are compared at different epigenetic marks and transcriptional factor binding sites and in different genome functions.

Consistent with the existing conclusions,^31,32^ we observe that nucleosome positioning levels at epigenetic marks and transcriptional factor binding sites correlate more with location rather than the effect of the epigenetic binding (Fig. 3A). In general, modified nucleosomes at promoters tend to have higher positioning levels, followed by enhancers and gene bodies. Within a particular region, the positioning level at activation modification is slightly higher than repression modification, i.e., H3K9ac and H3K4me2 compared with H3K9me3 and H3K27me3 in promoter regions. Besides histone modifications, histone variants and all tested chromatin-binding proteins also associate with higher positioning levels. For example, nucleosomes at structural proteins such as CTCF and RAD21 binding sites have higher positioning levels than the genome-wide average level. A consistent trend is confirmed by the nucleosome positioning levels in different chromatin states annotated by ChromHMM (Fig. 3B). In promoters and regions with high transcription activities, nucleosomes are better positioned, whereas in enhancers and repressed regions such as polycomb repressed regions and heterochromatin, they are more poorly positioned.

**Figure 3:**
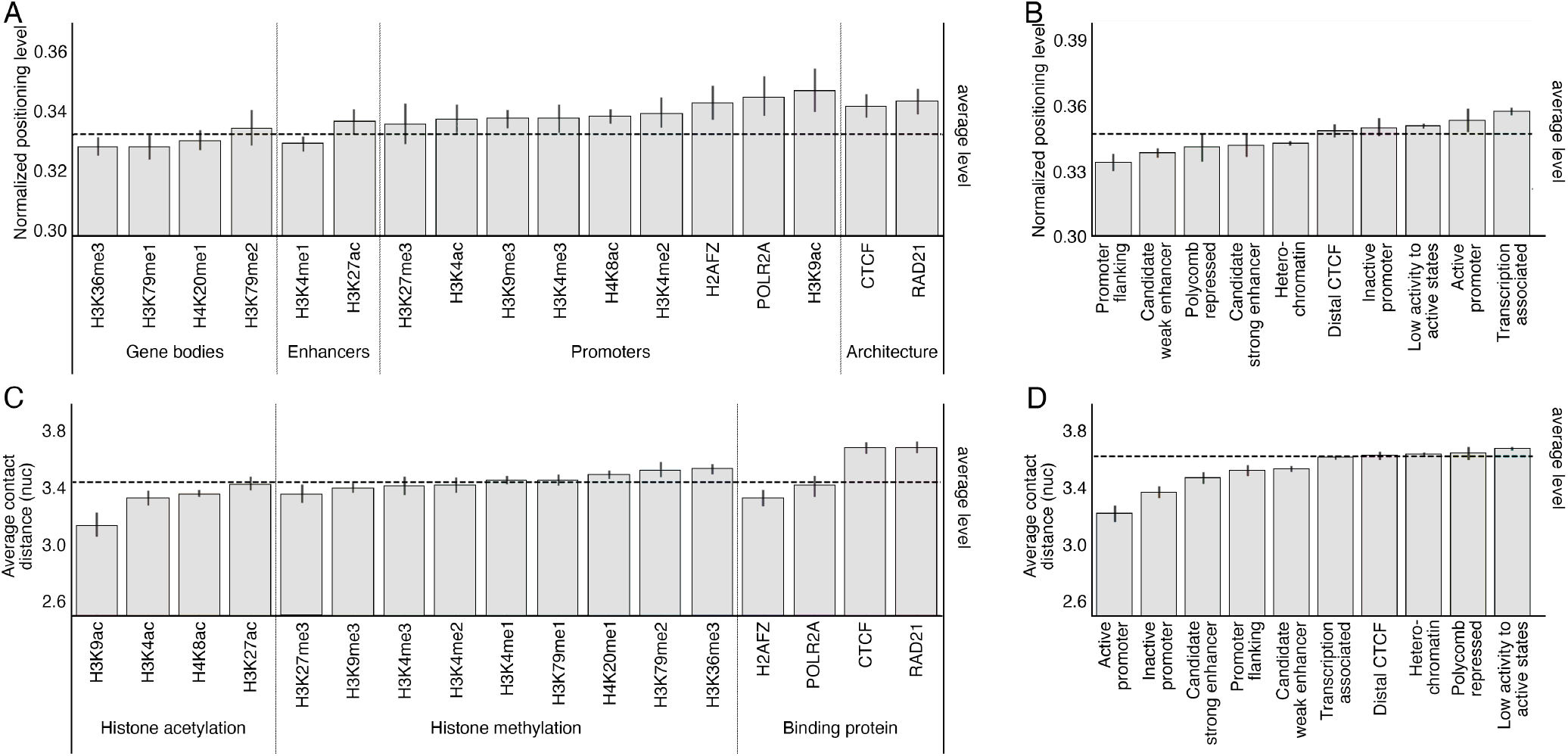
Nucleosome positioning levels and spatial organization stratify patterns of epigenetic modifications and genome functions. **A.** Normalized nucleosome positioning level at nucleosomes subject to specific epigenetic modifications or protein bindings. Generally, nucleosomes modified by promoter enriched epigenetic marks or at chromatin architecture associated protein binding sites are better positioned. Meanwhile, nucleosomes modified by gene body enriched or enhancer enriched epigenetic marks are more poorly positioned. **B.** Normalized nucleosome positioning level at nucleosomes within different chromatin states predicted by ChromHMM. Similar to the observation in epigenetic modifications, nucleosomes in promoters are better positioned, while nucleosomes in promoters and enhancers are more poorly positioned. Transcription associated state represents loci of RNA polymerase binding or mRNA elongation, which mostly occur near the active promoter. **C.** Average contact distance at nucleosomes subject to specific epigenetic modifications or protein bindings. A longer contact distance suggests more compact nucleosome spatial organization, while a shorter contact distance suggests more relaxed nucleosome spatial organization. Generally, nucleosomes modified by histone methylations are more tightly packed than nucleosomes modified by histone acetylations, but the spatial organization of nucleosomes at protein binding sites varies across different protein functions. **D.** Average contact distance at nucleosomes within different chromatin states predicted by ChromHMM. Compared with other states, nucleosomes at enhancers and promoters are more lightly packed regardless of their activities, resulting in shorter average contact distances.

In order to investigate the association between local spatial nucleosome distribution and genome functions, we calculate the average short-range contact distances (mean of contact distances which are ≤ 1kb) at different loci. Compared with histone methylation, histone acetylation tends to correlate with a shorter average contact distance, indicating a more relaxed chromatin fiber structure (Fig. 3C). This result is consistent with the fact that histone acetylation is enriched at euchromatin where the chromatin fiber is lightly packed. Meanwhile, the influences of binding proteins on nucleosome spatial organization are various. Long average contact distances are observed at structural protein binding sites such as CTCF and RAD21, consistent with the previous studies that these proteins mediate chromatin looping and other structures.^33^ Short average contact distances are observed at transcription-associated protein POLR2A and histone variant H2AFZ. These factors are enriched at active TSSs, consistent with the fact that euchromatin having relaxed structures are enriched with expressed genes.^34,35^ In addition, nucleosome spatial organization is also correlated with chromatin states and transcription activities (Fig. 3D). Short average contact distances are observed in promoters, enhancers, and the promoter flanking regions, indicating the chromatin is loose in these regions. Furthermore, average contact distances at active promoters are shorter than inactive ones, and strong enhancers are shorter than weak ones, suggesting that the nucleosomes are more lightly packed in regions more associated with transcription events.

### 3.4 Tetra-nucleosome folding motifs closely correlate with genome functions and chromatin structures

Two types of tetra-nucleosome folding structures, *α*-tetrahedron and *β*-rhombus, have been discovered in yeast and human by electronic microscopes in the previous studies.^26, 36,37^ Although these structures are modeled in ultra-high resolution contact maps in yeast,^26^ they are not yet modeled in human cell lines. To investigate the distribution of these structures, we predict the folding structures in hESC using machine learning models trained on the yeast genome.^26^

Since the spatial distance between two nucleosomes is inversely proportional to some constant order of the contact frequency,^38^ we extract 4×4 submatrices along the diagonal of the contact matrix as the input features. Ideally, chromatin contacts in the submatrices characterize the neighborhood of the nucleosomes. We observe in yeast that the number of contacts is closely related to the nucleosome folding structures (Table S1). In brief, *β*-rhombus tend to form neighborhoods with fewer contacts, while *α*-tetrahedron tend to form neighborhoods with more contacts. In order to improve the accuracy of prediction, we divide the features into four groups according to the proportions of *α*-tetrahedrons with respect to the contact numbers (Fig. S2). After that, ten commonly used classifiers are trained and compared in each group respectively (Table S2). At last, the models with the highest F1-scores in group2 (with 200-400 neighboring contacts), group3 (with 400-600 neighboring contacts), and group4 (with over 600 neighboring contacts) are selected and applied to hESC. Due to the overall low F1-scores, folding motifs of nucleosomes in group1 (with less than 200 neighboring contacts) are not predicted.

The ratio of predicted *α*-tetrahedron and **β**-rhombus in human (51.4% vs. 48.6%) are consistent with that in yeast (50.9% vs. 49.1%). We also observe that contact patterns in the neighborhood of *α*-tetrahedron and *β*-rhombus in human are similar to the patterns in yeast (Fig. S3). Together, these results imply that the classifiers trained on yeast successfully distinguish the folding motifs in human.

To investigate the correlation between the structural preference and genome function, we first compare the ratio of *α*-tetrahedron to *β*-rhombus at epigenetic marks, transcriptional factor binding sites, and candidate cis-regulatory element (cCRE) annotations. Surprisingly, almost all selected epigenetic marks and transcriptional factors exhibit a preference towards *α*-tetrahedron at their binding sites. On the contrary, the preferences on folding motifs vary among cis-regulatory elements. Three of the four cis-regulatory elements have certain preferences on the folding motifs. Higher levels of *α*-tetrahedron are observed at distal enhancers and enhancers, and more *β*-rhombus are observed at insulators. At promoters, the proportions of *α*-tetrahedron and *β*-rhombus are close to their global levels (1). Combining the results from epigenetic marks, transcriptional factor binding sites and cis-regulatory elements, it implies that although epigenetic marks and transcriptional factors bindings have a preference towards *α*-tetrahedrons, high-order genome functions still influence the final preference on tetra-nucleosome folding motifs.

Meanwhile, the distribution of folding motifs also highly correlates with large-scale chromatin structures. We observe different proportions of *α*-tetrahedron and *β*-rhombus at multiple chromatin structures including compartments, topologically associated domain (TAD) boundaries, stripes, and loops. *α*-tetrahedrons present more frequently in compartment A (expression-active chromatin), while higher proportion of *β*-rhombuses is observed in compartment B (expression-inactive chromatin) (1). At the level of nuclear subcompartments revealed by SPIN,^39^ we also observe consistent results. Among the eight identified SPIN-states, the highest proportions of *α*-tetrahedrons are found at two active states “Interior Active 1” (58.5%) and “Speckle” (58.6%), whereas the lowest proportions of *α*-tetrahedrons are found at inactive states “Lamina” (37.6%) and “Near Lamina 1” (37.6%) (1). Moreover, we find that the preference on folding motifs changes at TAD boundaries according to their strength. Rigid (“strong”) boundaries tend to form more *β*-rhombuses, and permissive (“weak”) boundaries tend to form more *α*-tetrahedrons (1). Previous studies have shown that the strength of TAD boundaries is associated with their functionalities,^40^ possibly explaining the difference in their preferences on folding motifs. With respect to loops and stripes, higher proportions of *α*-tetrahedrons are observed (Table 1). One possible explanation of the preference towards *α*-tetrahedron in these regions is that compacted local domains in chromatin contact maps, such as loop extrusion, play a role in the formation of compartment A.^41^

**Table 1:** Preferences on tetra-nucleosome folding motifs in different regions.

| Regions | Prop. of $\alpha$ motif | Prop. of $\beta$ motif | Folding motif ratio | Preference | Significance |
| --- | --- | --- | --- | --- | --- |
| hESC chr21 | 51.4% | 48.6% | 1.00 | NA | ns |
| **Epigenetic marks and transcriptional factor binding sites** |  |  |  |  |  |
| CTCF | 51.5% | 48.5% | 1.00 | NA | ns |
| H3K27ac | 51.9% | 48.1% | 1.02 | α-tetrahedron | ** |
| H3K36me3 | 51.7% | 48.3% | 1.01 | α-tetrahedron | * |
| H3K4me1 | 53.2% | 46.8% | 1.12 | α-tetrahedron | **** |
| H3K4me2 | 52.6% | 47.4% | 1.05 | α-tetrahedron | **** |
| H3K4me3 | 51.8% | 48.2% | 1.02 | α-tetrahedron | *** |
| H3K79me2 | 51.6% | 48.4% | 1.00 | NA | ns |
| H3K9ac | 51.7% | 48.3% | 1.01 | α-tetrahedron | ** |
| H3K9me3 | 51.7% | 48.3% | 1.01 | α-tetrahedron | * |
| H3K18ac | 51.6% | 48.4% | 1.00 | NA | ns |
| Nanog | 51.9% | 48.1% | 1.02 | α-tetrahedron | ** |
| Rad21 | 51.6% | 48.4% | 1.00 | NA | ns |
| H2AFZ | 52.1% | 47.9% | 1.03 | α-tetrahedron | ** |
| GTF2F1 | 51.6% | 48.4% | 1.00 | NA | ns |
| **cis-Regulatory elements** |  |  |  |  |  |
| Distal enhancers | 52.2% | 47.8% | 1.03 | α-tetrahedron | **** |
| Enhancers | 54.5% | 45.5% | 1.14 | α-tetrahedron | ** |
| Promoters | 50.9% | 49.1% | 0.98 | β-rhombus | *** |
| Insulators | 46.3% | 53.7% | 0.813 | β-rhombus | *** |
| **Chromatin compartments** |  |  |  |  |  |
| Compartment A | 54.6% | 45.4% | 1.14 | α-tetrahedron | **** |
| Compartment B | 46.8% | 53.2% | 0.83 | β-rhombus | **** |
| **SPIN states** |  |  |  |  |  |
| Interior active1 | 58.5% | 41.5% | 1.34 | α-tetrahedron | **** |
| Interior active2 | 49.2% | 50.8% | 0.92 | β-rhombus | **** |
| Interior active3 | 43.9% | 56.1% | 0.74 | β-rhombus | **** |
| Interior repressive2 | 46.3% | 53.7% | 0.81 | β-rhombus | **** |
| Lamina | 37.6% | 62.4% | 0.57 | β-rhombus | **** |
| Near lamina1 | 41.7% | 58.3% | 0.67 | β-rhombus | **** |
| Near lamina2 | 42.3% | 57.7% | 0.69 | β-rhombus | **** |
| Speckle | 58.6% | 41.4% | 1.35 | α-tetrahedron | **** |
| **TAD boundaries** |  |  |  |  |  |
| Strong boundaries | 47.8% | 25.2% | 0.86 | β-rhombus | **** |
| Weak boundaries | 69.2% | 30.8% | 2.12 | α-tetrahedron | **** |
| **Other chromatin structures** |  |  |  |  |  |
| Loops | 57.5% | 42.5% | 1.27 | α-tetrahedron | **** |
| Stripes | 55.4% | 44.6% | 1.17 | α-tetrahedron | **** |

Folding motif ratio is calculated by comparing the $\alpha/\beta$ ratios in specific regions and the genome-wide $\alpha/\beta$ ratio. A folding motif ratio greater than 1 indicates that the region has a preference towards $\alpha$ -tetrahedrons, and a folding motif ratio smaller than 1 indicates a preference towards $\beta$ -rhombus.

### 3.5 Nucleosome contact maps provide enhanced details

Traditionally, chromatin contact maps are generated at a fixed resolution, e.g. 5kb or 10kb. Since we now reach the nucleosome resolution, we hypothesize that a nucleosome contact map is more appropriate and informative to visualize their 3D organization.

Unlike traditional bin-based contact maps in which the nodes represent fixed-length genome fractions, nodes in nucleosome contact maps represent nucleosomes. Reads assigned to the different nucleosomes constitute the contacts between different nodes, illustrating the spatial proximity between these nucleosomes.

Compared with 200bp-resolution contact maps, nucleosome contact maps contain more interpretable and equivalently precise contact patterns. While having similar numbers of N/N+1, N/N+2, N/N+3, and N/N+4 contacts as 200bp-resolution contact maps, nucleosome contact maps barely include self contacts (N/N contacts), suggesting that most contacts connect two different nodes in nucleosome contact maps (Fig. 4A). This property is consistent with the fact that every contact in the ultra-high resolution contact map consists of reads from two different nucleosomes. Although the contact distribution changes, nucleosome contact maps still achieve the same level of precision as the commonly used 200bp-resolution bin-based contact maps, measured by the distance error between the aligned reads and their assigned node centers (Fig. 4B). In addition, the error in nucleosome contact maps is symmetrically distributed compared with 200bp-resolution contact maps, because nucleosomes called by NucleoMap reflect the distribution of aligned reads, providing a more accurate presentation of the intrinsic inter-nucleosomal structures. Therefore, nucleosome contact maps capture the nucleosome organization within the nucleus more precisely than traditional bin-based chromatin contact maps.

**Figure 4:**
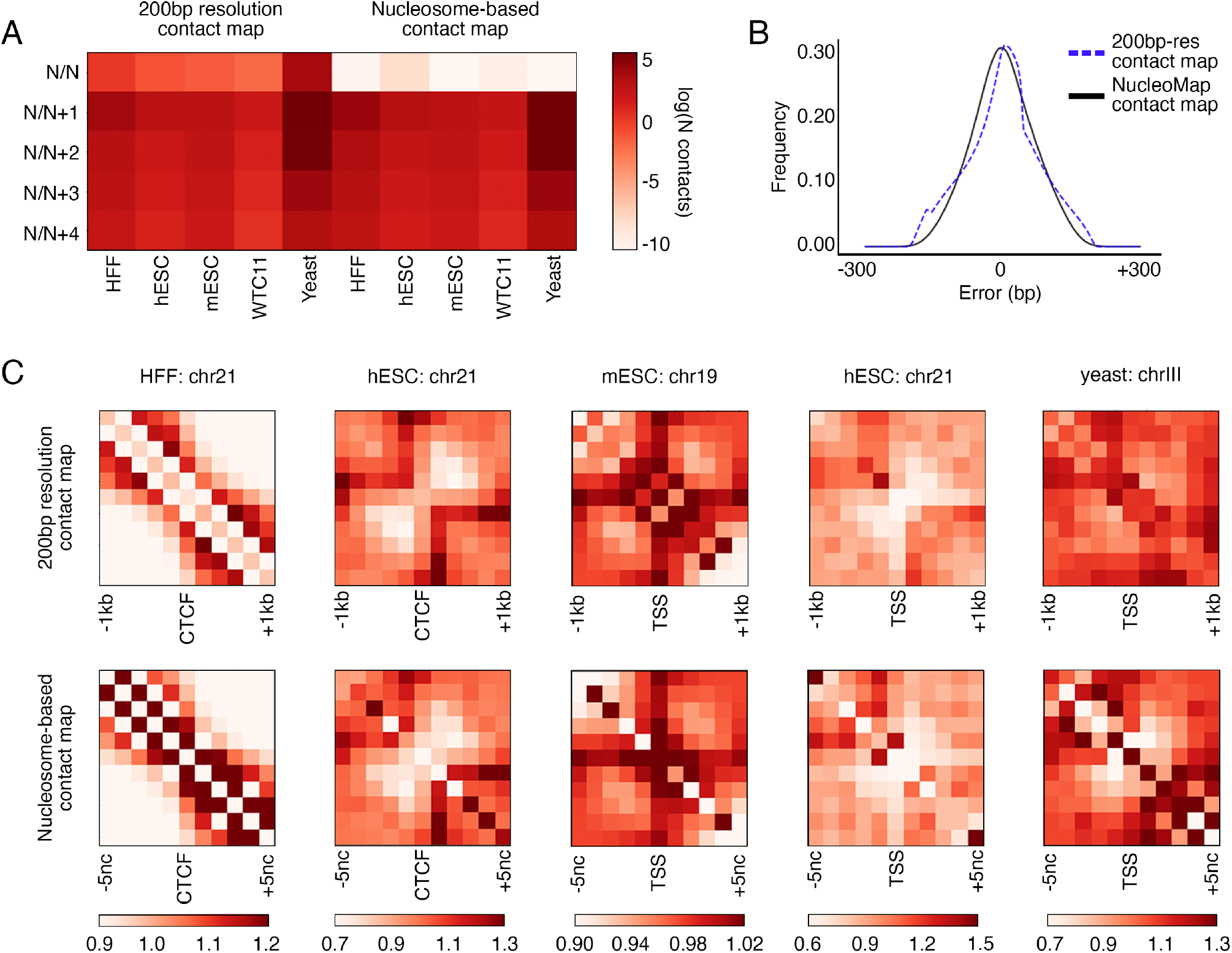
Nucleosome contact maps constructed by NucleoMap contain more concentrated inter-nucleosomal contact signals. **A.** Averaged contact numbers between neighboring nodes in 200bp-resolution contact maps and NucleoMap nucleosome contact maps. Compared with the 200bp-resolution contact maps, the number of self-contacts (N/N contacts) significantly decreases in nucleosome contact maps, which is more intuitive because two ends of a contact connect different nucleosomes in a cell. **B.** Frequencies of contact distance errors after assigned to 200bp bins (blue) and nucleosomes identified by NucleoMap (black). The distance error of every contact is defined as the difference in contact distance after assigning both ends to their corresponding nodes in a chromatin contact map. Nucleosome contact maps achieve the same level of precision as 200bp-resolution contact maps. **C.** OE Normalized pileup nucleosome contact maps and 200bp-resolution contact maps centered by CTCF binding sites or TSS regions in different cell lines. Nucleosome arrays are separated into two domains by CTCF binding sites and TSS flanking regions. Compared with 200bp-resolution contact maps, nucleosome contact maps reveal more concentrated patterns in most cell lines.

Compared with 200bp-resolution bin-based contact maps, nucleosome contact maps better recover fine-scale nucleosomal structures. Nodes in traditional 200bp-resolution contact maps may not accurately cover the DNA wrapping around nucleosomes, and thus interactions between mono-nucleosomes are not precisely captured by the contacts between nodes. In contrast, contacts between two nodes in nucleosome contact maps intuitively represent the proximity of two nucleosomes. Since nucleosomes are basic structural components of chromatin, the maps better illustrate the fine-scale nucleosomal motifs. In the pileup maps centered by CTCF binding sites and TSSs in five cell lines, nucleosome contact maps provide more concentrated signals (Fig. 4C). A stronger contrast between the low contact frequency background and the high contact frequency looping structures anchored at CTCFs or TSSs is shown in the nucleosome contact maps, allowing easier identification of spatial nucleosome motifs.

## 4 DISCUSSION

Incorporating inter-nucleosome distance information reveals more detailed and precise nucleosome positioning throughout the genome. Here we report a computational approach, NucleoMap, for nucleosome identification in both well-positioned and poorly positioned regions from ultra-high resolution contact maps. Using public Micro-C data from yeast, human, and mouse, we demonstrated that NucleoMap effectively detects nucleosomes in complex mammalian genomes, where most nucleosomes are poorly positioned. As the resolution of 3D chromatin organization profiling reaches the nucleosome level, nucleosome contact maps are more intuitive than classical fixed-bin resolution contact maps.

The genome-wide nucleosome positioning identified by NucleoMap provides an opportunity to revisit epigenetic mark data at mono-nucleosome resolution. Although it has long been known that epigenetic marks are decorated on mono-nucleosomes, previous studies rarely explore the mono-nucleosome level due to the ubiquitously distributed poorly positioned nucleosomes in complex eukaryote genomes. The genome-wide nucleosome map enhances existing epigenetic mark data to mono-nucleosome level, especially in the poorly positioned transcriptionally silent chromatin. Furthermore, by integrating epigenetic modifications and properties of nucleosome arrays in different genome regions, it is possible to establish a more comprehensive understanding of gene regulation. However, we note that more accurate mapping of epigenetic signals to mono-nucleosomes requires both ultra-high resolution epigenetic signals such as CUT&RUN data and enhanced computational approaches that consider the densities of mono-nucleosomes and epigenetic signals. Follow-up work is still required to design computational methods specifically for mono-nucleosome level sequencing data mapping.

The produced nucleosome contact maps allow a comprehensive analysis of the association between nucleosome spatial organization and genome functions. Although some computational approach has been developed to model inter-nucleosomal contacts,^42^ NucleoMap is the first method jointly identifying nucleosomes and modeling inter-nucleosomal contacts. Using the nucleosome contact map constructed by NucleoMap, the hierarchical chromatin structures such as tetra-nucleosome folding motifs are retrieved by *in silico* approaches in the human genome. It is possible to identify more accurate tetra-nucleosome folding motifs and higher-level chromatin folding structures with the help of enhanced machine learning models that utilize the sequential nature of the chromatin. Furthermore, combined with the epigenetic signals annotated to mono-nucleosomes, it is also possible to establish a 3D framework illustrating the spatial structures of epigenetic events. Compared with traditional studies in this area which focus on the interactions along the linear DNA sequence, this framework unveils interactions of chromatin modifications in an ultra-high resolution 3D space, and thus providing additional knowledge in the regulation of genome activities.

## 5 METHODS

### 5.1 NucleoMap algorithms

#### Estimating read centers in Micro-C contact maps

In order to estimate the true read density along the genome, we need to first estimate positions of read centers, which are not directly accessible from the contact map data. Each contact in chromatin contact maps is composed of two anchor reads, with various sizes from ~120bp to ~170bp, referring to different nucleosomes (Fig. S4 step 4). During the paired-end sequencing and downstream processing, only ~50bp at two ends of a contact, one for each read, are sequenced and mapped to the reference genome, thus the centers of both reads are not contained in the alignment data (Fig. S4 step 6). Therefore, read centers need to be estimated from the mapped read ends.

One effective way to estimate read centers is to shift the ends towards 3’ direction by half of the average read size. NucleoMap automatically estimates the average read size in the Micro-C data using the difference in contact distances across contact types. Four types of contacts, ++, +−, −+, and −−, can be found in the chromatin contact map data, according to the strands the reads mapped to (Fig. S5). Contacts of different types vary in contact distance even when they anchor the same nucleosome pair. +− contacts cover two nucleosome dyads and the fragment between them, and ++ contacts and −− contacts cover a nucleosome dyad and the fragment between them, while −+ contacts cover only the fragment between them. Based on this observation, the average read size is calculated as the average difference in contact distances between +− contacts and ++ contacts with three steps. First, NucleoMap calculates the contact distance distributions of short-range +− and ++ contacts. After that, peaks are called from the two distributions. The first peak centers in the histograms correspond to the average contact distance between neighboring nucleosomes in +− and ++ contacts. Finally, the average read size is estimated to be the distance between the first peak centers in the two distributions.

After shifting reads to their centers, both reads in the contacts with genomic distances shorter than 160bp are excluded in the downstream analysis to prevent artifacts introduced by the outliers.

#### Calculating sequence-based binding score

Sequence-based binding score measures sequence-based nucleosome affinity at a given position. The score is calculated by normalizing the convolution score of an AA/AT/TA/TT dinucleotide PWM.

The binding score is calculated in four steps. First, NucleoMap calculates a Position Frequency Matrix (PFM) of AA/AT/TA/TT dinucleotides. PFM records the occurrences of AA/AT/TA/TT dinucleotides at each position within ±100bp from the *N* read centers. Based on the dinucleotides frequency, a 2 × 200 PFM *F* is generated by

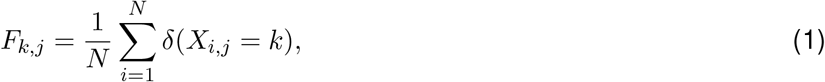

where *i* ∈ [1, *N*], *j* ∈ [1, 200] and *k* = {0, 1} indicates the occurrence of AA/AT/TA/TT dinucleotides. *δ* is an indicator function. The first row of the matrix indicates the occurring frequency of AA/AT/TA/TT dinucleotides, and the second row indicates the occurring frequency of other dinucleotides. Following that, a PWM *W* expressing the binding patterns is calculated as

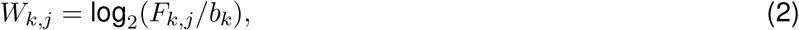

where *b_k_* denotes the background frequencies of AA/AT/TA/TT dinucleotides and other dinucleotides calculated from the reference genome. In the third step, NucleoMap calculates the cross correlation scores *D* between the PWM and the one-hot encoded reference genome.

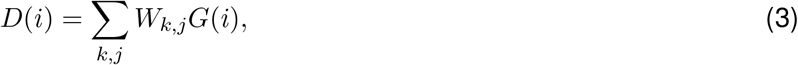

where *G*(*i*) is the dinucleotide in reference genome at position *i.* This score illustrates the similarity between the nucleosome binding pattern and the genome sequence at a given position. In the last step, a binding score *B* is generated by normalizing the cross correlation score *D*. The cross correlation score is first normalized over its neighborhood,

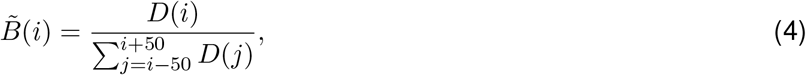

and then *z*-normalized,

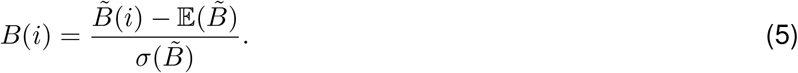

The resulting binding score represents the magnitude of increase in nucleosome binding preference at a position compared with its neighborhood.

#### Estimating nucleosome numbers and defining the objective function

Reads are assigned to nucleosomes within 1kb using a hard clustering DP mixture model with a fixed covariance *σI*. We assume that within 1kb on the genome, reads *X* = {*x_i_*} are samples drawn from an unknown number of Gaussian distributions with fixed covariance *σ*, representing the dyad of a nucleosome. Under this assumption, a DP mixture model of nucleosomes is formulated as follows:

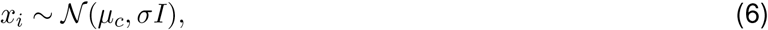

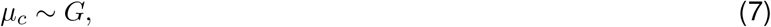

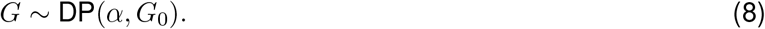

Here 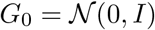 is a prior over the mean distributions of the Gaussian mixtures, and a draw 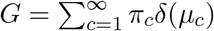 is the mean distribution of a Gaussian mixture, where *π*_c_ denotes the weight of the *c*-th Gaussian. For *i* = 1, 2,…, *n*, the probability *p_c_* of assigning a read *x_i_* to an existing nucleosome *c* is

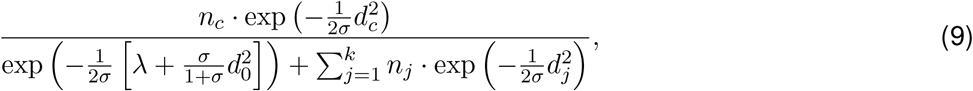

where *n_c_* is the number of reads assigned to nucleosome *c*, 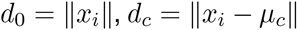, and 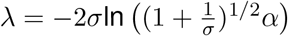. Similarly, the read *x_i_* is assigned to a new nucleosome with a probability *p_new_*

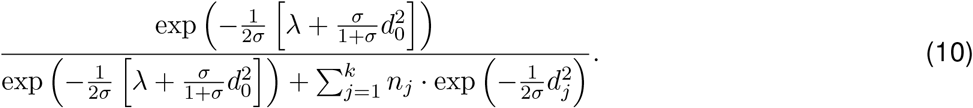

A hard assignment DP mixture model is obtained by pushing *σ* → 0. When *σ* approaches 0, the numerator of *p_new_* is dominated by *λ*. Furthermore, as *σ* → 0, the assignment probabilities become binary and only the smallest values of 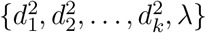 receive a non-zero probability. In particular, a new nucleosome is created whenever a read is farther than 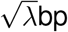 away from every existing nucleosome center. The underlying objective of this model is similar to the *k*-means objective function,

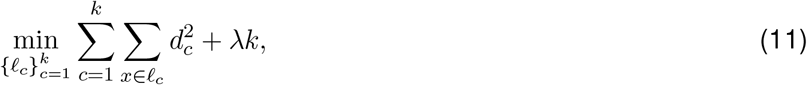

where *ℓ_c_* is the set of reads assigned to nucleosome *c*. The threshold *λ* controls the trade-off between the traditional *k*-means term and the cluster penalty term. Optimizing this objective function identifies potential nucleosome centers based on the read density.

#### Integrating read density, pairing information, and binding scores

To incorporate pairing information and binding scores, an adjusted distance 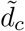 is used instead of *d_c_.* We define

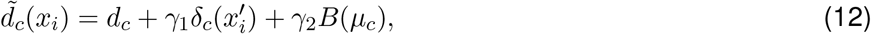

where 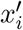 is the other read sharing a contact with *x_i_, δ_c_* an indicator function returning 1 if 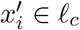 and 0 otherwise, and *γ*_1_, *γ*_2_ the corresponding distance penalties. The final objective function is

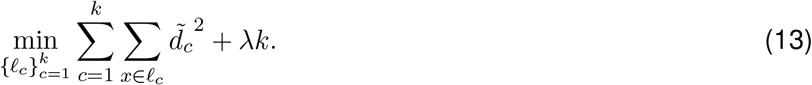

The model is optimized using a previously proposed hard clustering algorithm that behaves similarly to *k*-means with the exception that new clusters are formed when the aforementioned condition is satisfied.^43^

### 5.2 Identifying nucleosomes from Micro-C data

Alignment files of Micro-C data are downloaded from 4DN data portal (human cell lines), or generated by Bowtie2 with ‘very sensitive’ mode (mESC and yeast) using reference genomes hg38, mm10, and SacCer3 respectively.^44^ Mapped reads from all replicates are merged before calling nucleosomes. Using the alignment files and the following parameters, we benchmarked NucleoMap and multiple baseline methods including DANPOS2,^15^ NOrMAL,^16^ Nseq,^18^ and NucTools.^20^ NucleoMap is run with default parameters. DANPOS2 is run with parameters “-m 0 -p 0.05”. NOrMAL is run using the original config.txt on its GitHub repository. Nseq is run with parameters “-f 0.01 -s 10 -t 16”. NucTools is run with parameters “-dir -use -w 100”. Since the output of NucTools is an occupancy profile, nucleosome centers are called by the peak calling function signal.find peaks in python scipy package with the parameter “distance=160”. For DANPOS2 and NucTools, the alignment files are first transformed into .bed format. For all baseline methods, we select the single-end MNase-seq mode and ignore the pairing relationships within a contact.

### 5.3 Calculating nucleosome occupancy

Nucleosome occupancy measures the fraction of nucleosomes covering a given position in a cell population. The measure is originally proposed by Valouev et al.^14^ to describe the nucleosome positioning level, but here a smaller neighborhood *w* = 30 is chosen in the normalization step to increase its detection sensitivity.

The nucleosome occupancy is calculated in three steps. First, a smoothing kernel *K* is defined as

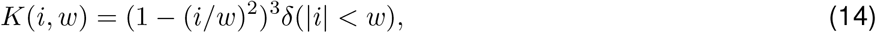

where *w* defines an aggregation window and *δ* is an indicator function. In the second step, we generate the read coverage files from the alignment files using samtools depth with parameters “-a -H -Q 10”.^45^ After that, the convolution kernel *K* is applied to the read coverage file along the chromatin

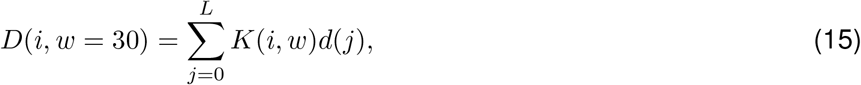

where *L* is the length of the chromatin and *d*(*j*) represents the number of read centers at position *j*. At last, the smoothed density is normalized over its neighborhood

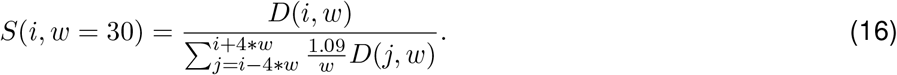

A scaling factor 1.09 is designed to normalize the occupancy values as

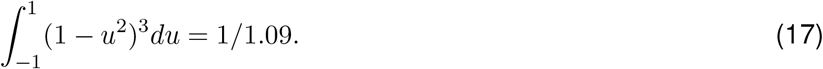

The neighborhood size in the denominator is set to ±4 * *w* such that it covers a slightly larger region than a well-positioned nucleosome (146bp) to capture the poorly positioned nucleosomes.

### 5.4 Annotating genome features to mono-nucleosomes

Epigenetic modification peaks are assigned to the nearest nucleosomes to the peak centers. In this way, we generate binarized signals indicating whether or not a nucleosome is subjected to certain modifications, and peak strengths and fold changes are ignored. Similarly, mono-nucleosome positioning levels are calculated using the nucleosome occupancy signal. We define the highest occupancy value within ± 30bp from a nucleosome center as its normalized positioning level. Cis-regulatory elements are annotated to all nucleosomes within a ± 500bp neighborhood.

Nucleosomes within the span of compartments, SPIN states, or stripes are assigned with the corresponding features. TAD boundaries are annotated to all nucleosomes within a ± 500bp neighborhood. Loops are annotated to all nucleosomes within a ± 500bp neighborhood at each anchor.

### 5.5 Predicting tetra-nucleosome folding motifs

For the *i*-th nucleosome, we generate a 10-dimension feature using elements from the upper triangle of the sub-contact-matrix containing the (*i* − 1)-th, the *i*-th, the (*i* + 1)-th, and the (*i* + 2)-th nucleosomes. Tetra-nucleosome motif labels of the yeast genome are collected from the nucleosome 3D coordinates generated in a published study.^26^

Nucleosomes are grouped according to the sum of their features. Ten classifiers from the sklearn python package are trained in each group, including k-Nearest Neighbors, Linear SVM, RBF SVM, Gaussian Process, Decision Tree, Random Forest, Multilayer Perceptron, AdaBoost, Gaussian Naive Bayes, and Quadratic Discriminant Analysis. The parameters in these models are as follow: KNeighborsClassifier(k=3), SVC(kernel=“linear”, C=0.025), SVC(gamma=2, C=1), GaussianProcessClassifier(1.0 * RBF(1.0)), DecisionTreeClassifier(max depth=5), RandomForestClassifier(max depth=5, n estimators=10, max features=1), MLPClassifier(alpha=1, max iter=1000), AdaBoostClassifier(), GaussianNB(), and QuadraticDiscriminantAnalysis().

In each group, 75% of the nucleosomes are randomly selected as training data, and the others are used as test set. The classifiers are trained on the training data, and their performances are evaluated by F1-scores on the test sets.

The folding motif preference is measured by a folding motif ratio within a specific region, defined as

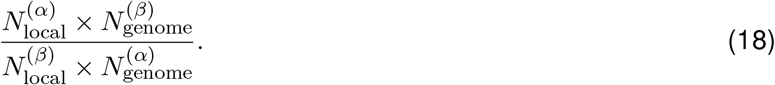

When this ratio > 1, the region has a preference towards *α*-tetrahedron and towards **β**-rhombus otherwise. The significance of folding motif preferences is evaluated using two-tail T-tests. Folding motif ratios are compared between nucleosomes within specific regions and nucleosomes sampled from the whole genome.

### 5.6 Constructing nucleosome contact maps and OE normalization

Nucleosome contact maps are constructed by assigning contacts to their corresponding nucleosomes identified by NucleoMap.

NucleoMap estimates the expected contact numbers between nucleosome pairs according to their genomic distance. Based on the assumption that the contact frequency is a function of genomic distance, the expected contact numbers between two nucleosomes is estimated given their genomic distance *d*,

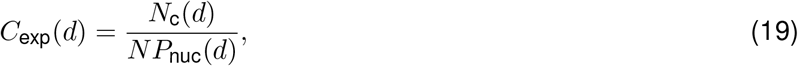

where *N_c_* refers to the total number of contacts in the contact map with genomic distance *d*, and *NP*_nuc_ refers to the total number of nucleosome pairs in the contact map with genomic distance *d*.

However, it is difficult to calculate *NP*_nuc_ directly in practice because it requires a computational complexity of O(*n*^2^), where *n* is the number of nucleosomes in the contact map. To avoid the expensive computation, we instead estimate *NP*_nuc_ with a summation over multiple Erlang distributions. Assuming that nucleosomes occur at a steady rate along the genome, the genomic distances between neighboring nucleosomes follow an exponential distribution

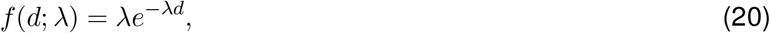

where *d* is the genomic distance and *λ* = *L*_chrom_/*N*_nuc_ is the occurring rate of nucleosomes. Therefore, the genomic distance between the *i*-th and the (*i* + *k*)-th nucleosomes follow an Erlang distribution which characterizes the sum of *k* independent exponential distributions

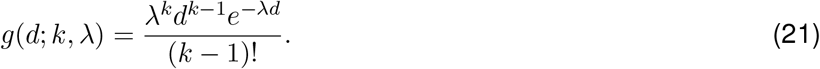

Hence the probability of having *k* nucleosomes within a certain range of genomic distance [*d*_1_, *d*_2_], denoted by *P*(*d*_1_, *d*_2_; *k, λ*), is calculated by the difference in CDF of the Erlang distribution,

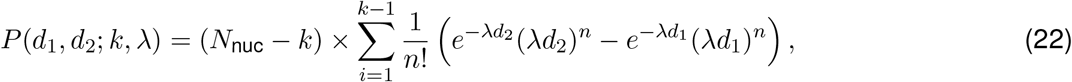

and the expected number of nucleosome pairs within the range [*d*_1_, *d*_2_] in the chromatin, denoted by *NP*_nuc_(*d*_1_, *d*_2_), is calculated by summing the differences of multiple Erlang distributions under a series of *k*s,

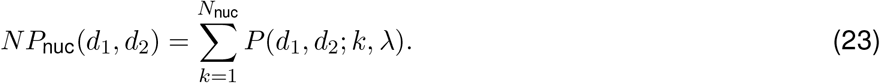

To further reduce computational complexity, this number is approximated by a smaller set of *k*s

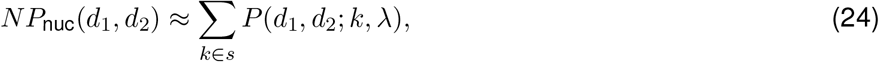

where max(1, *d*_1_/150 – 20) ≤ *s* ≤ min(*N*_nuc_, *d*_2_/150 + 20). The OE normalized contacts between two nucleosomes are finally given by the ratio between observed contacts and the expected contacts,

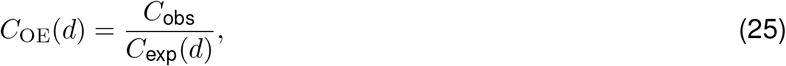

where *C*_obs_ is the contact numbers between the nucleosome pairs, and *d* is their genomic distance.

## 6 DATA ACCESS

Majority of the sequencing data involved in the this paper are public available in NCBI GEO repository,^46^ ENCODE project,^13^ and 4DN data portal.^47^ All software in this paper are available on GitHub. A python implementation of NucleoMap is provided on GitHub, which takes processed contact pair files as input and generates nucleosome-based contact maps (see Table 2).

**Table 2:**
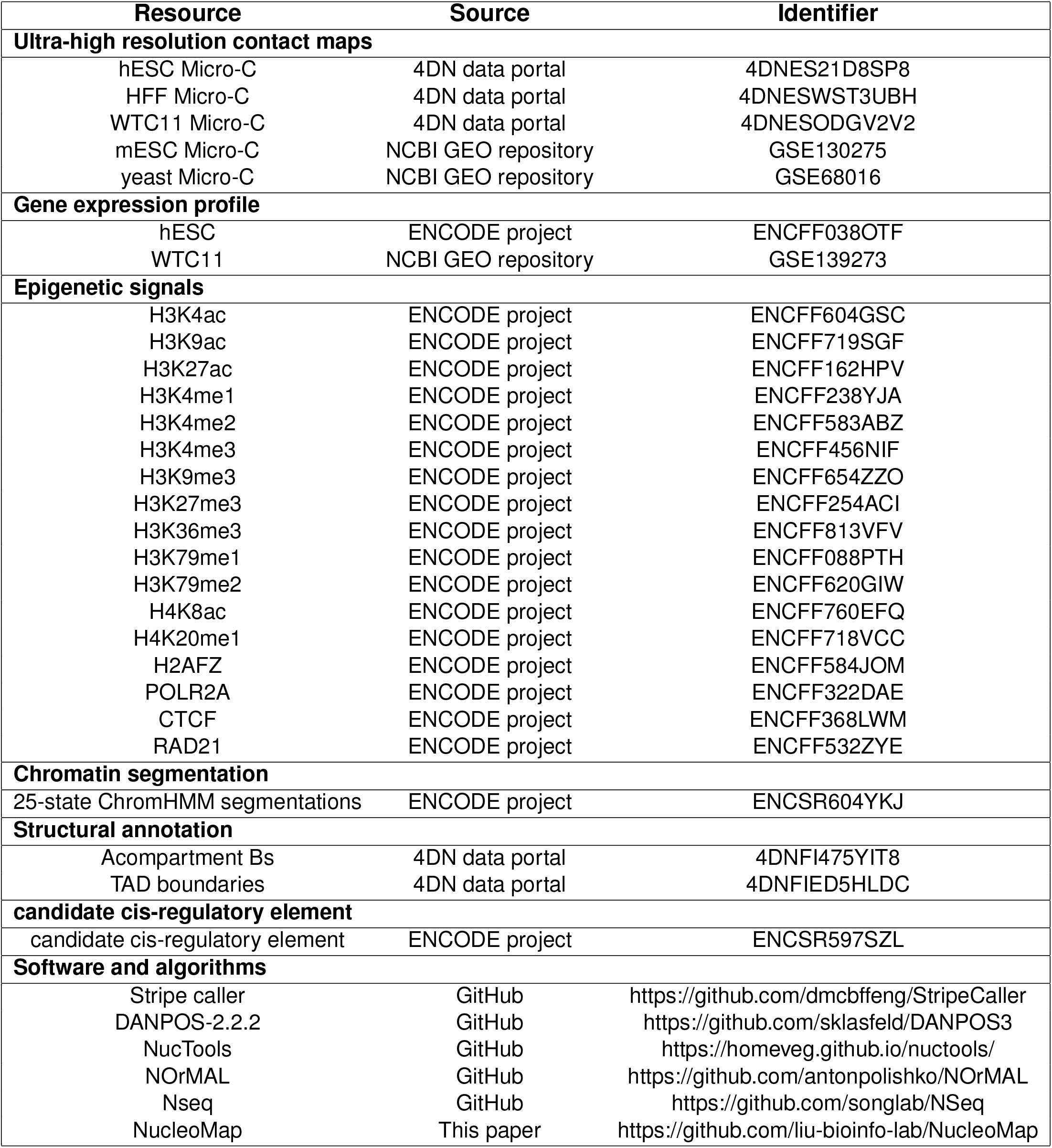
Data and softwares involved in this paper.

Loops in hESC micro-C data are called with Juicer HiCCUPS algorithm.^48^ Stripes in hESC micro-C data are called by the stripe caller developed by our group (see Table 2). SPIN state data are generated in a published study.^39^

## 8 ACKNOWLEDGEMENTS

This research is funded by NIH award R35HG011279.

